# A sea full of measures: EU conservation goals for benthic habitats will require wide-ranging spatial measures

**DOI:** 10.64898/2026.05.11.724278

**Authors:** Wolfgang Nikolaus Probst

**Affiliations:** Thuenen-Institute of Sea fisheries, Herwigstraße 31, 27572 Bremerhaven, Germany

**Keywords:** Marine Strategy Framework Directive, Nature Restoration Regulation, marine benthic habitats, fisheries closures, marine protected areas, offshore renewables, spatial optimisation

## Abstract

The use of marine space by human activities is globally increasing, resulting in a competition with spatial management measures for marine conservation. Within the European Union (EU) these measures are currently implemented by the union member states to achieve the UN sustainable development goal (SDG) of protecting at least 10 % of the national marine waters. Further, the EU Marine Strategy Framework Directive (MSFD) and the Nature Restoration Regulation (NRL) are the two main legal means for the implementation of ambitious spatial conservation targets for benthic habitat types, which can range from 10 – 90 %. This study analysis how the targets of the MSFD and NRL are currently met in the German waters of the North Sea and which areas the full implementation of both legislations might require. A spatial optimisation tool (“prioritizr” in R) was used to identify optimised solutions for the conservation of up to 75 % of NRL benthic habitats. The current spatial conservation measures (which ban demersal trawling within certain zones of designated marine protected areas, MPA) are not sufficient to reach the targets of the MSFD and NRL. Extending the exclusion of demersal trawling to the entire area of the MPAs would achieve a sufficient coverage for all habitats except for offshore sand and mud habitats. These could be further protected, when including areas for offshore wind farms, where trawling is also banned. However, to date it is unclear, if and how these (or other human use) areas could be included into spatial conservation regimes, a debate that needs to be resolved to allow for the achievement of the ambitious MSFD and NRL targets.

## Introduction

The human use of marine space has drastically intensified in the course of the 20^th^ century (Watson et al., 2012; Halpern et al., 2025). In the North Sea, this global development can be seen as under a magnifying glass (Emeis et al., 2015; Piet et al., 2023), as an increasing number of marine activities such as shipping, fishing, construction of marine infrastructure such as offshore renewables and cables and pipelines, extraction of oi, gas, sand and gravel, military defence, or carbon capture and storage require marine space (Burrows et al., 2021; OSPAR, 2023; Stelzenmüller et al., 2024). As a result of the intensified human activities, marine spatial planning (MSP) has emerged as a predominant mean for governmental authorities to regulate marine uses in space and time (Frazão Santos et al., 2019; Trümpler et al., 2023). And at the same time, spatial measures (SPM) have become a highly relevant tool for marine conservation restricting human use in designated areas (often referred to as marine protected areas or marine reserves, hereafter refererred to as MPA; Day et al., 2019; Kriegl et al., 2021). The establishment of MPA as gained momentum at the end of the 0^th^ century globally (Maestro et al., 2019; Sala et al., 2021), as well as in the North Atlantic (Mazaris et al., 2019; Roessger et al., 2022; EEA, 2024).

In 2019 the European Commission (EC) presented the Green Deal, an environmental policy strategy that combines aspects of climate protection, waste reduction and biodiversity conservation (Paleari, 2022). For the latter, the latest policy released by the EC in 2024 was the EU Nature Restoration Law (NRL), which obliges EU member states to restore and protect marine habitats within their national waters (Hering et al., 2023; Bodenbender, 2024). The NRL thereby complements and aligns with previous EU directives and policies, i.e. the Marine Strategy Framework Directive (MSFD), the Habitats Directive (HD), the Birds Directive (BF), the Water Framework Directive (WFD), the Common Fisheries Policy (CFP) and the EU Biodiversity strategy 2030. All of these directives and polices include the obligation to implement MPAs for the conservation of benthic habitats, marine birds & mammals or sensitive life stages of commercially relevant fish stocks. As a consequence, member states of the European Union have designated and implemented an increasing number of MPAs in their national waters (EEA, 2024).

In article 5 of the NRL spatial restoration targets are defined for a wide range of marine habitat groups (see Table 1), i.e. 30 % recovery by 2030, 60 % recovery by 2040 and 90 % recovery by 2050 for each group, that did not achieve a good environmental status (GES). The restoration and protection of degraded habitats can be achieved by active (e.g. planting sea grass seedlings) or passive measures (e.g. banning bottom trawling), which member stats are supposed to present in a national restoration plan by 2026. As a result of the NRL and aforementioned EU policies, the political incentive (or pressure) for national governments to extent their network of spatial conservation measures is high.

**Table 1.** Relationship between NRL and MSFD habitat groups and types in German waters of the North Sea.

| NRL Habitat group | MSFD Habitat type |
| --- | --- |
| Seagrass beds | Littoral sediments [BHT], Plates [OHT] |
| Macroalgae forests | Infralittoral rock and biogenic reef [BHT] |
| Shellfish beds | Circalittoral coarse sediment [BHT] |
|  | Infralittoral coarse sediment [BHT] |
|  | Offshore circalittoral coarse sediment [BHT] |
|  | Mixed coarse gravel and sand [OHT] |
|  | Reef [OHT] |
| Maerl beds | Not present in German waters |
| Sponge, coral & coralligenous beds | Circalittoral coarse sediment [BHT] |
|  | Infralittoral coarse sediment [BHT] |
|  | Offshore circalittoral coarse sediment [BHT] |
|  | Mixed coarse gravel and sand [OHT] |
|  | Reef [OHT] |
| Vents & seeps | Not present in German waters |
| Soft sediments | Circalittoral mixed sediment [BHT] |
|  | Infralittoral mixed sediment [BHT] |
|  | Offshore circalittoral mixed sediment [BHT] |
|  | Circalittoral sand [BHT] |
|  | Infralittoral sand [BHT] |
|  | Offshore circalittoral sand [BHT] |
|  | Circalittoral mud [BHT] |
|  | Infralittoral mud [BHT] |
|  | Offshore circalittoral mud [BHT] |
|  | OSPAR mud habitats [OHT] |
|  | Sandbank [OHT] |

Benthic habitats are the main object of Article 5 of the NRL and their definition is provided in Annex II. These definitions are aligned with the nomenclature of the HD and the European nature information system (EUNIS) and are congruent with the definition of broad habitats types (BHT) of the MSFD. Apart from BHT, the MSFD also defines other habitat types (OHT), which are habitats that require particular protection under the HD or regional seas conventions such as the Oslo-Helsinki Commission (OSPAR). OHT thereby can be reefs, sandbanks, shallow bays with sea grass meadows, ocean vents, sponge and coral communities or muddy habitats with deep burrowing megafauna (EEC, 1992; Gutow et al., 2020).

The NRL aims at very high rates of coverage (up to 90 % until 2050), which poses the question, how marine space in general and sites for offshore renewables in particular could be designed to accommodate different forms of use and conservation (Stelzenmüller et al., 2016; Püts et al., 2023). The question of multi-use or multi-purpose designation of offshore windfarms (OWF) has initiated the discussion of whether these could be designated as ‘other effective area-based conservation measures’ (OECM), which until today is not yet resolved (EC, 2022; FAO, 2022). While some argue that the environmental impacts of OWF are too substantial to allow them to be considered as areas for conservation (Claudet et al., 2022; NABU, 2023), others argue that OWFs can support the recovery of sensitive species such as elasmobranchs (Hermans et al., 2025), enhancement of commercial fish and shellfish stocks (Stelzenmüller et al., 2021; Gimpel et al., 2023; Thatcher et al., 2023) and the increase and conservation of biodiversity (Knorrn et al., 2024; Dannheim et al., 2025). With increasingly limited marine space available, OWF might therefore need to be considered as complementary areas for conservation (Campbell et al., 2025). Accordingly, spatial management measures (SPM) might not be confined to MPA, but might also occur in areas of co-use such as OWF.

Demersal trawl fisheries have been demonstrated to be the main threat to the ecosystem status of benthic habitats (Rijnsdorp et al., 2020; Pitcher et al., 2022) and hence their exclusion can be considered as the measure of choice to restore degrade habitats (Hiddink et al., 2017; OSPAR, 2023). Accordingly, this study analyses the coverage of NRL habitat groups and MSFD habitat types by various spatial measures, i.e. implemented closures to demersal trawl fisheries (FCL), marine protected areas without fisheries exclusions (MPA) and offshore windfarms (OWF), which are or will become closed for fishing until 2040 (Stelzenmüller et al., 2022; Bonsu et al., 2024). To evaluate the possibilities for the achievement of NRL and MSFD targets, spatially optimized solutions in different spatial scenarios (e.g. by including FCL, MAP or OWF) were identified using the conservation optimization tool ‘prioritizr’ within the R-programming environment. The national German waters of the North Sea were considered as a case study for the implementation of the NRL to exemplify the high level of ambition and resulting, considering that the situation in the German waters of the North Sea is representative of many member states of the European Union.

## Material & methods

### Study site

The German waters of the North Sea include the costal sea (up to 12 nm offshore) and the exclusive economic zone (EEZ, Figure 1). Within the EEZ, the federal government implemented three MPAs (Borkum Reef, Dogger Bank & Sylter Outer Reef) with five fisheries exlcusion zones for demersal trawling.

**Figure 1.**
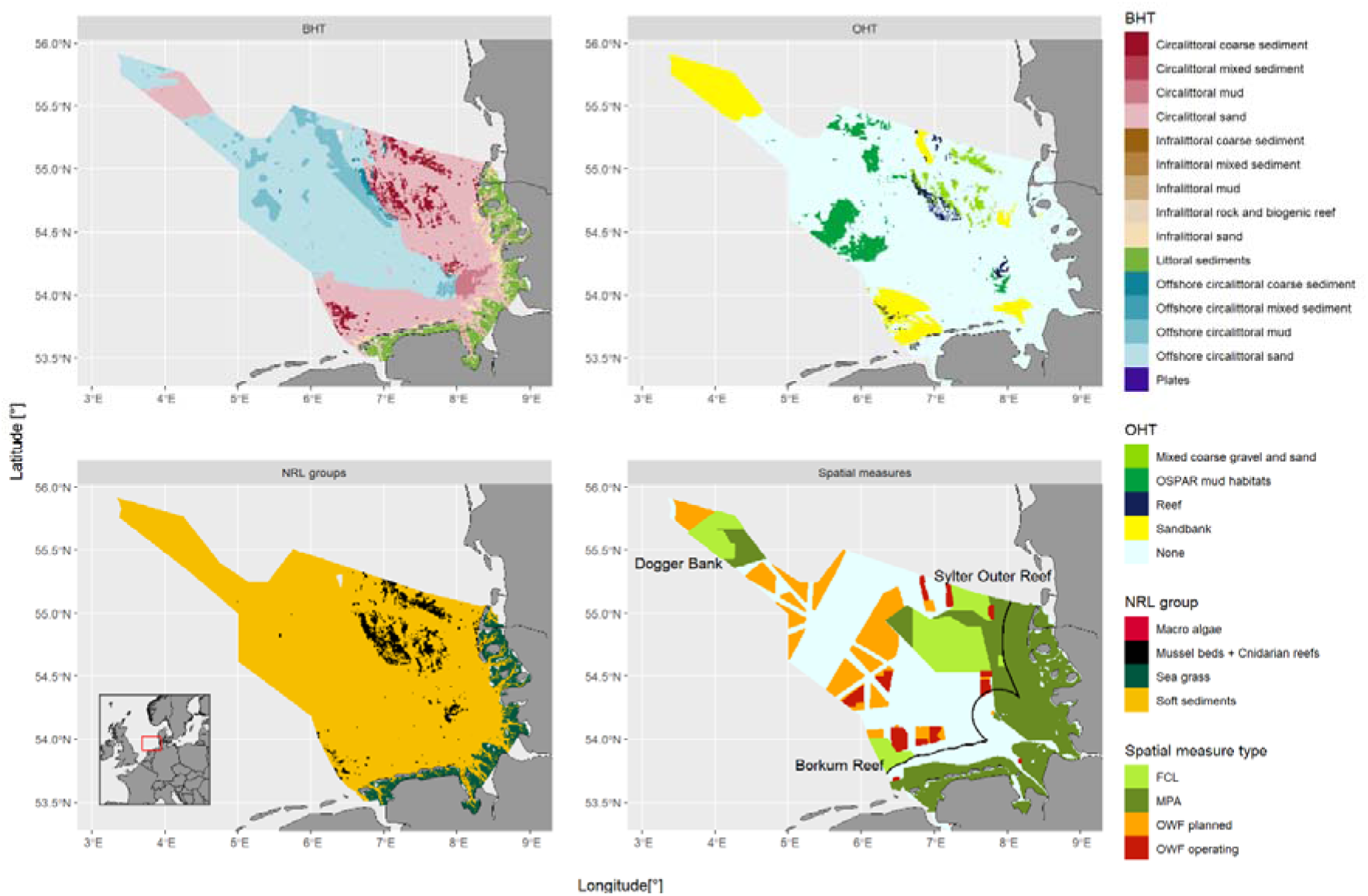
Location of study site, habitat types (of the MSFD) and habitat groups (of the NRL) and spatial management measures affecting fisheries access in the German waters of the North Sea. FCL=Closures to demersal trawl fisheries, MPA=Marine protected area (without any exclusion), OWF = offshore windfarm site.

Sites for offshore windfarms have been designated within the EEZ, with more than 1.200 of ∼ 8.700 planned turbines are already operating (as of 2024). To reduce the complexitiy of the the analysis, operating and planned windfarms were merged, assuming that existing and planned OWF sites will be developped until 2050. Further, OWF sites hereafter are included in the abreviation SPM; even though they are not a conservation measure by definition. However, they include a conservation aspect for benthic habitats, as they – depneding on country – might or will exclude demersal trawl fisheries.

### Data sources

Data of MSFD habitat types (BHT & OHT) were obtained from the German Federal Agency of Conservation (BfN) and merged to combine both habitat types into one map. Accordingly, all OHT-raster cells that were not NA were set to overwrite values in the according raster cell of the BHT-raster, thereby replacing the BHT-value. There was also a small section without available habitat data in the central northern part of the study area, which was excluded from the analysis.

The habitat groups of the NRL were derived from the MSFD habitat types by assigning these types to the NRL habitat groups (Table 1). It should be noted that MSFD habitat types such as reefs and coarse sediments were considered as suitable habitats for NRL groups 3 and 5, as both groups (mussel/oysters and cnidarians/sponges/coralligenous algae) were associated with reefs and coarse sediments as settling substrate. As to date, this approach is the official approach by the German authorities for determining the habitat extent of NRL groups 3 and 5. Further, NRL goup 2 (macro algae forests) were only considered to be present around the rocky substrate around the island of Heligoland, accounting for only 0.816 km^2^ (< 0.002 % of the study area of 41.290 km^2^). In all other areas the water is too turbid or too deep.

Shapefiles on fisheries closures (FCL) within the MPA were obtained from the BfN (Borkum Reef Ground and Sylter Outer Reef) and based on vertices as defined by EU/2017/118 (Dogger Bank).

Data on offshore windfarm sites were obtained from 4COffshore Ltd (last valid year was 2024). The sites were filtered for the statuses “Concept/Early Planning”, “Consent Authorised”, “Development Zone”, “Fully Commissioned”, “Partial Generation/Under Construction”, “Pre-Construction”, “Under Construction” to include operating and planned windfarms while excluding cancelled, failed and dormant projects. OWF shape files were buffered by 500 m to account for the safety zone around OWF, in which fishing is not permitted (Bonsu et al., 2024).

Data on the German maritime spatial plan 2021 were obtained from geodata information site of the Federal Maritime and Hydrographic Agency (https://gdi.bsh.de/de/data/Maritime-Spatial-Plan-in-the-German-Maritime-Area.zip).

All shapefiles were rasterized on a grid of 0.0025° longitude x 0.0025° latitude resulting in grid cells with an average cell area of 0.045 km^2^ (min = 0.044 km^2^, max = 0.046 km^2^).

### Overlap analysis of current and future SPM

The spatial optimisation software package ‘prioritizr’ for the statistical computing language R was used for all subsequent analysis.

To determine the spatial overlap of FCL, MPA and existing/planned OWF with MSFD habitat types and NRL groups, ‘prioritizr’ was configured with a minimum target of zero percent coverage the according areas (FCL, MPA including FCL, OWF & MPA + OWF) as locked-in-constraints, leading the algorithm to determine the overlap (i.e. coverage) between spatial measures (SPM) and the habitat types/groups (for the R code see supplements S1). The total Ferrier-importance (FI) was calculated for as sum of Ferrier-importance for each habitat type/group to identify the most relevant areas within each SPM. Ferrier-importance is a measure of irreplaceability i.e. how relevant a given grid cell is for the achieved ‘prioritizr’-solution (Ferrier et al., 2000).

### Scenarios

The second objective of this study was to determine the potential requirements of additional SPM when the spatial objectives of MSFD/NRL were to be met until 2050. These spatial targets were defined as shown in Table 2 for each MSFD habitat type and NRL habitat group. For the NRL habitat Group 7 – soft sediments, German authorities currently set a restoration target of 75% by 2050, which deviates from the 90 % target for the other habitat groups.

**Table 2.** Habitat type/group targets for 15 modelling scenarios (3 variations of NRL targets x 5 combinations of spatial measures [SPM]).

| Scenario # | SPM | BHT target | OHT target | NRL targets3 |  |
| --- | --- | --- | --- | --- | --- |
|  |  |  |  | NRL target<br>[Groups 1, 2, 3 & 5 –<br>sea grass,<br>macro algae,<br>mussel beds<br>& sponge/cnidarian reefs] | NRL<br>[Group 7 –<br>Soft<br>sediments] |
| 1 | No constraints | 0.1 | 0.9 | 0.3 | 0.3 |
| 2 | FCL | 0.1 | 0.9 | 0.3 | 0.3 |
| 3 | FCL + MPA | 0.1 | 0.9 | 0.3 | 0.3 |
| 4 | OWF | 0.1 | 0.9 | 0.3 | 0.3 |
| 5 | FCL + MPA + OWF | 0.1 | 0.9 | 0.3 | 0.3 |
| 6 | No constraints | 0.1 | 0.9 | 0.6 | 0.6 |
| 7 | FCL | 0.1 | 0.9 | 0.6 | 0.6 |
| 8 | FCL + MPA | 0.1 | 0.9 | 0.6 | 0.6 |
| 9 | OWF | 0.1 | 0.9 | 0.6 | 0.6 |
| 10 | FCL + MPA + OWF | 0.1 | 0.9 | 0.6 | 0.6 |
| 11 | No constraints | 0.1 | 0.9 | 0.9 | 0.75 |
| 12 | FCL | 0.1 | 0.9 | 0.9 | 0.75 |
| 13 | FCL + MPA | 0.1 | 0.9 | 0.9 | 0.75 |
| 14 | OWF | 0.1 | 0.9 | 0.9 | 0.75 |
| 15 | FCL + MPA + OWF | 0.1 | 0.9 | 0.9 | 0.75 |

‘Prioritizr’ was set to search for minimal areas, which accommodated these targets, again with FCL, MPA, OWF and MPA + OWF as locked-in constraints (supplements S2). The applied penalty (= 0.01) and edge factor (=0.5) were validated empirically to find a combination that produces comprehensive SPM networks (high representativeness of habitat types/groups with low boundary length and area costs) while reducing computational effort. To further reduce the computational load, raster data on feature and cost matrices were re-scaled by a factor of ten resulting in a grid resolution of 0.025 ° longitude x 0.025 ° latitude and the time limit for the cbc-solver was set to 1.800 seconds.

## Results

### Overlap of existing and future SPM

FCL covered only 8 out of 15 BHT, but three out four OHT, and two out of four NRL-groups (Figure 2). Including MPA areas increased the coverage for all habitats except for offshore circalittoral sand and mud [BHT], OSPAR mud habitats [OHT] and soft sediments [NRL]. These gaps however, could be closed by OWF areas.

**Figure 2.**
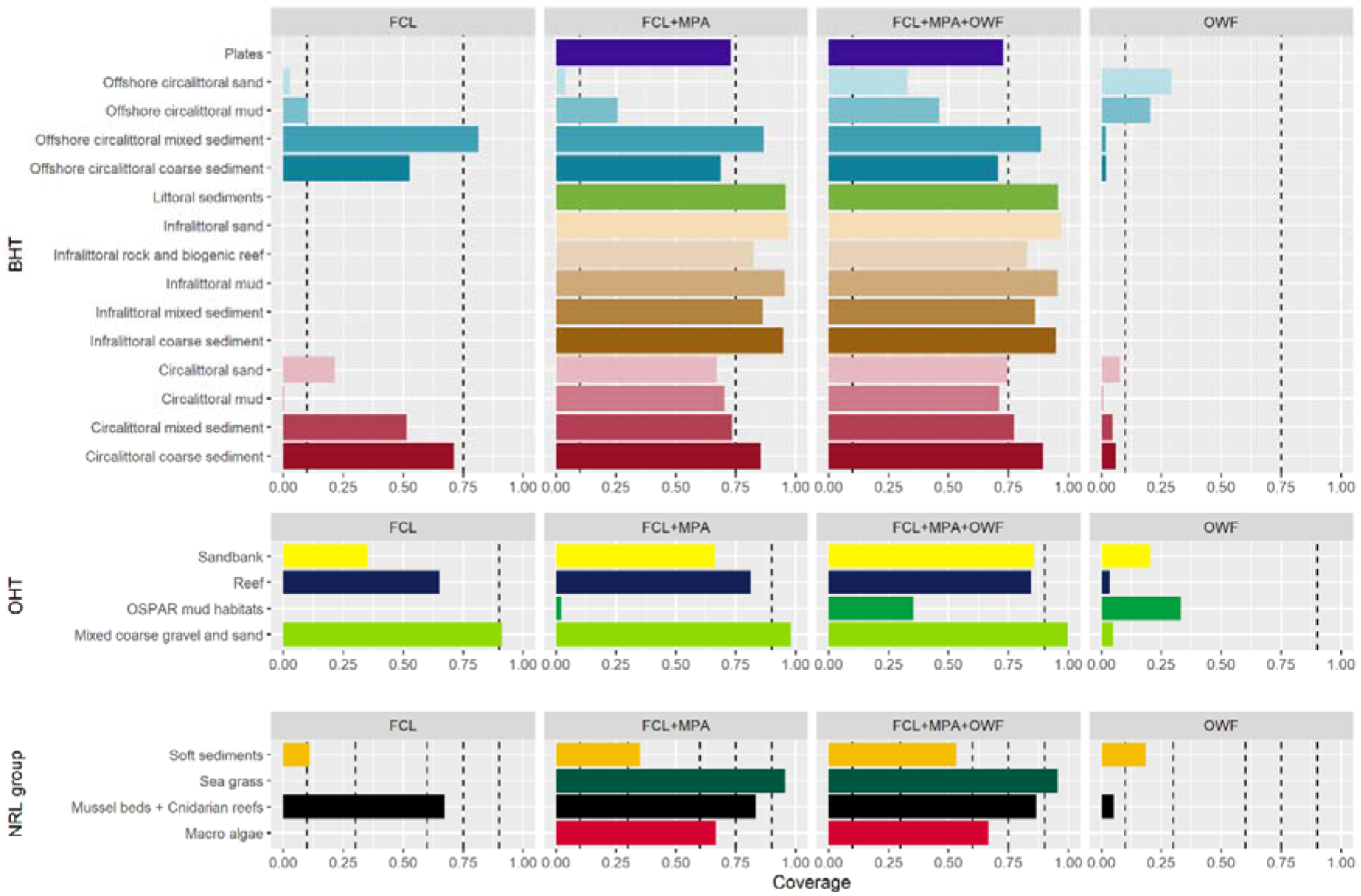
Relative coverages of MSFD habitat types and NRL habitat groups within different spatial measures. FCL = closures to demersal trawling, MPA = marine protected areas including FCL, OWF = offshore wind farms, MPA + OWF = combined areas of MPA including FCL and OWF. Dashed lines represent spatial objectives of the national German MSFD/NRL. The 0.75-line represents potential German national targets for NRL soft sediment habitats and MSFD BHT (to be further evaluated).

The mean relative coverages of the different SPM indicated that MPA (including FCL) could provide an average coverage of at least 60 % for all habitat types (BHT, OHT, NRL; Table 3). By including OWF, this coverage could be increased to at least 75 %, with OWF contributing the highest average coverage to OSPAR mud habitats.

**Table 3.**
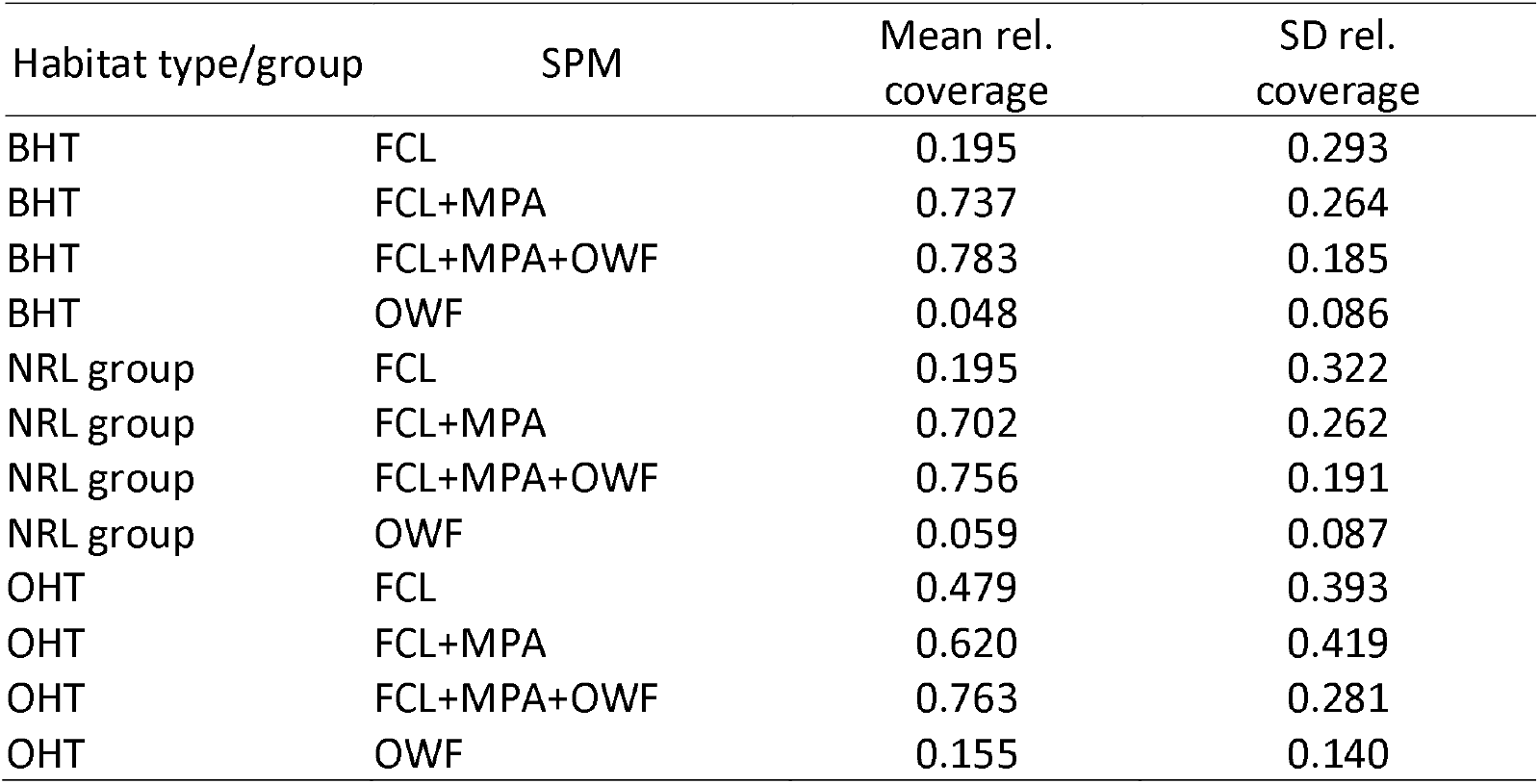
Mean relative coverage of habitat types/groups under existing spatial measures (SPM).

### Ferrier importance (FI) of covered areas

FI increased with increasing area of SPMs, with maximum values being ∼0.3 when considering only FCL and increasing to 0.78 when including FCL, MPA and OWF (Figure 3). Mussel beds and circalittoral coarse sediments (being both OHT and NRL) achieved the highest FI in all SPM scenarios.

**Figure 3.**
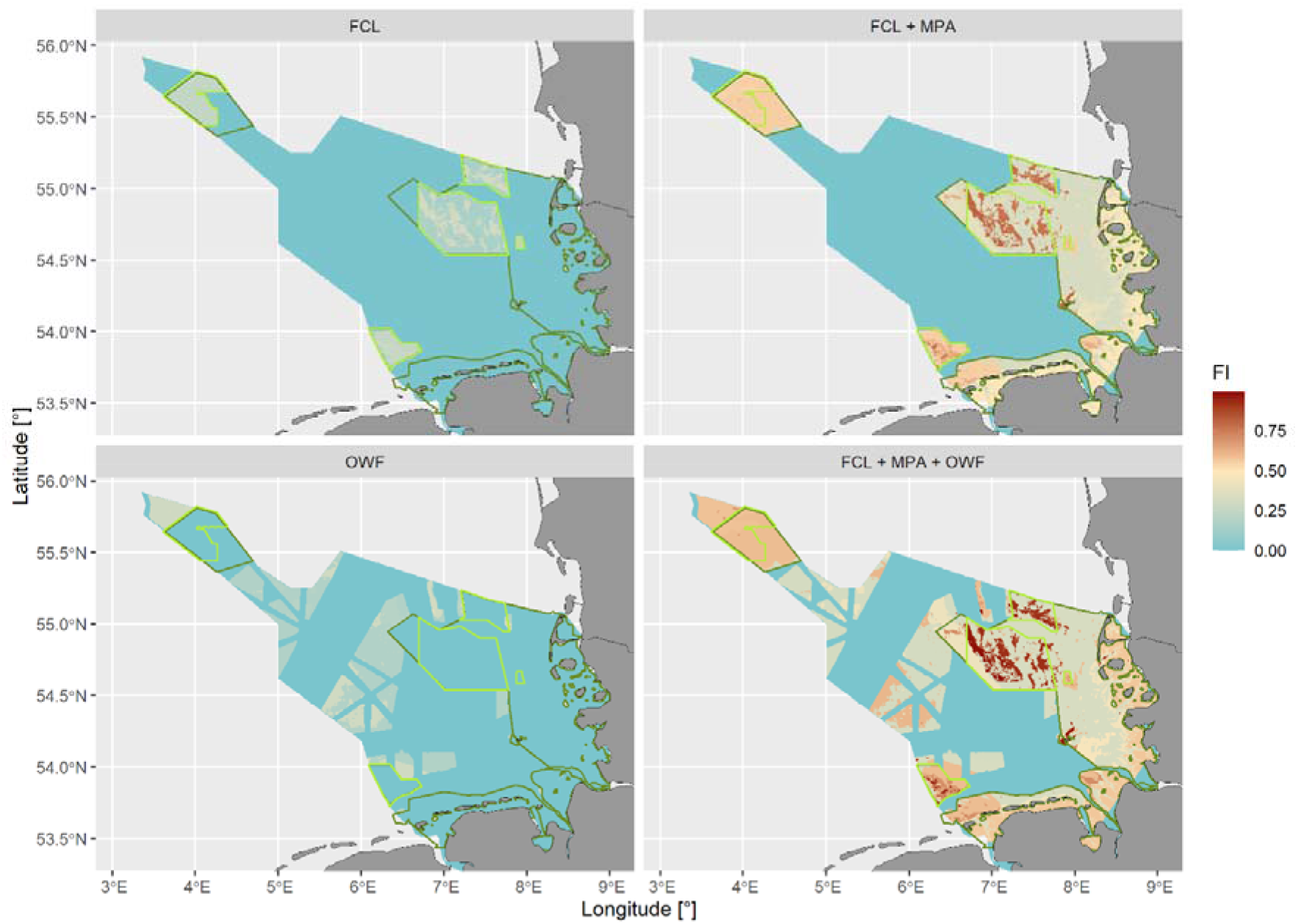
Ferrier-importance (FI) for the different coverages of spatial management measures. Abbreviations as in Figure 1.

### Scenarios of SPM

The hypothetical scenario without locked-in constraints (i.e. without any existing SPM), shows the lowest spatial demands with the 30-%-NRL-coverage target (Figures 4 & 5). Interestingly, in with this NRL-target the second lowest areal demands were achieved by the OWF scenario with requiring less area outside any SPM. With the 60 %- and 75/90 %-NRL-targets, the un-constrained solutions did not require less area than the scenarios with SPM. Instead, the least solution area outside any SPM was required with the FCL+MPA+OWF scenarios.

**Figure 4.**
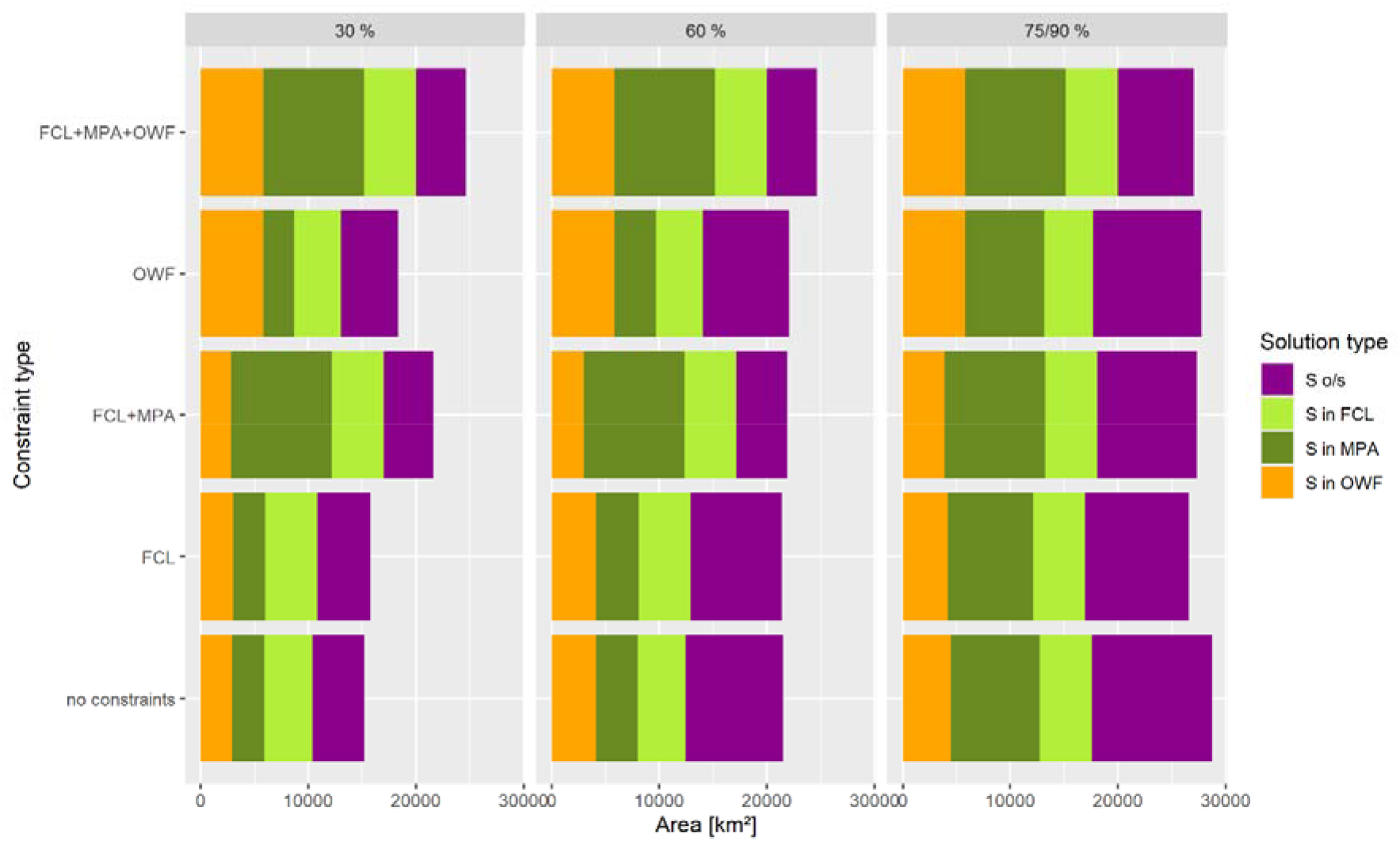
Area requirements for different spatial conservation targets of the MSFD and NRL. Constraint type refers to using spatial measures as locked-in contraints in the ‘prioritizr’-solutions.

In the 75/90 %-scenarios the overall areal demand of each ‘prioritizr’-solution was ∼30.000 km^2^, amounting to around ∼73 % of the study area, whereas in the 30%- and 60%-scenarios the areal demand achieved these target values.

## Discussion

The results of this study clearly demonstrate that the simultaneous achievement of the MSFD & NRL benthic habitat targets will result in conservation and restoration measures covering ∼ 75 % of the national German waters of the North Sea by 2050. And even though many measures in these areas might rather be passive than active measures, this still might drastically limit the space for maritime activities which cause physical disturbances on the seabed. The “prioritizr”-solutions for the maximum conservation targets (scenarios 11 – 15) overlap substantially with areas designated for human activities (military defence, construction and of cables, sand and gravel extraction, scientific research including experimental trawling, construction of offshore windfarms and fishing) as designated in the German marine spatial plan from 2021 (Trümpler et al., 2023; Figure 6). The full implementation of the MSFD and NRL spatial conservation targets for benthic habitats might thereby result in spatial conflicts with almost all marine human activities. Amongst these, fisheries might be the most impacted, as this activity is currently the most spread and considered as particularly detrimental to the ecological status of benthic habitats (Kaiser et al., 1998; Gerritsen et al., 2013; Pitcher et al., 2022).

The German fishing sector is in a state of transformation as it is confronted with ecological and political challenges (Lasner and Barz, 2023). Many traditional stocks like North Sea cod *Gadus morhua* or herring *Clupea harengus* have become scarce, while the access to traditional fishing grounds has become restricted, either due to Brexit (Harte et al., 2019) or the construction of OWF (Stelzenmüller et al., 2022). The full implementation of the MSFD and NRL targets might further force the transformation of the German fishing sector from benthic trawling towards less invasive (i.e. passive) gears such as crab pots (Stelzenmüller et al., 2021; Barz et al., 2025) and new resources such as cephalopods (Sulanke et al., 2025).

A ban on benthic trawling on the full area of the MPA network would allow to cover at least 60 % of all benthic habitat types but offshore sand and mud habitats (see Figure 2). These could be complemented by OWF areas to achieve a coverage of 25 – 45 %. However, until to date it is debated whether the installation of OWF would have detrimental or beneficial impacts on benthic habitats (NABU, 2023; Dannheim et al., 2025). While the foundations of wind turbines can increase biodiversity by attracting sessile benthic species and their consumers (Krone et al., 2013; Degraer et al., 2020; Dannheim et al., 2025), their construction causes disturbances and loss of original benthic habitats (Watson et al., 2024). Further impacts on benthic habitats might result from laying cables, ship traffic and noise. The protective benefits of trawling bans on benthic habitats in and around OWF might therefore be counterbalanced by detrimental impacts during the construction, operation and decommissioning (Watson et al., 2024). To allow for the inclusion of OWF into spatial conservation measures, it could be argued that only a fraction of the total OWF area should be considered against spatial conservation goals of the original habitats. For example, an OWF with an area of 25 km^2^ which is established on a sandy offshore habitat, might only add 10 km^2^ to the spatial conservation target for this habitat type, as the remaining 15 km^2^ might be affected by foundations and cables (note that this is an arbitrary example, the exact fraction would depend on the number of turbines, the length of set cables and the estimated spatial footprint of turbines, foundations and cables).

This study has a national focus analysing the situation within the German waters in the southern North Sea, which can be considered as representative for the situation in many other member states of the European Union (e.g. Belgium, the Netherlands, Denmark, Sweden or Poland), which are confronted with the national implementation of the same policy obligations from various EU directives and regulations, and at the same time, with similar spatial limitations. The ambitious goals of the NRL, particularly the 75 %/90 %-goal by 2050 might therefore bear the same conflict potential between human activities and nature conservation in other countries than Germany. Further, conflicts between MSP and spatial conservation measures might also arise outside of the EU in countries where the NRL does not apply. Nonetheless, many nations have committed themselves to implement the sustainable development goals (SDG) of protecting 30 % of their nationals marine waters, of which 10 % should be strictly protected (UN, 2015). Figure 5 exemplifies that even these spatial conservation targets may cause conflicts between marine conservation and human activities.

**Figure 5.**
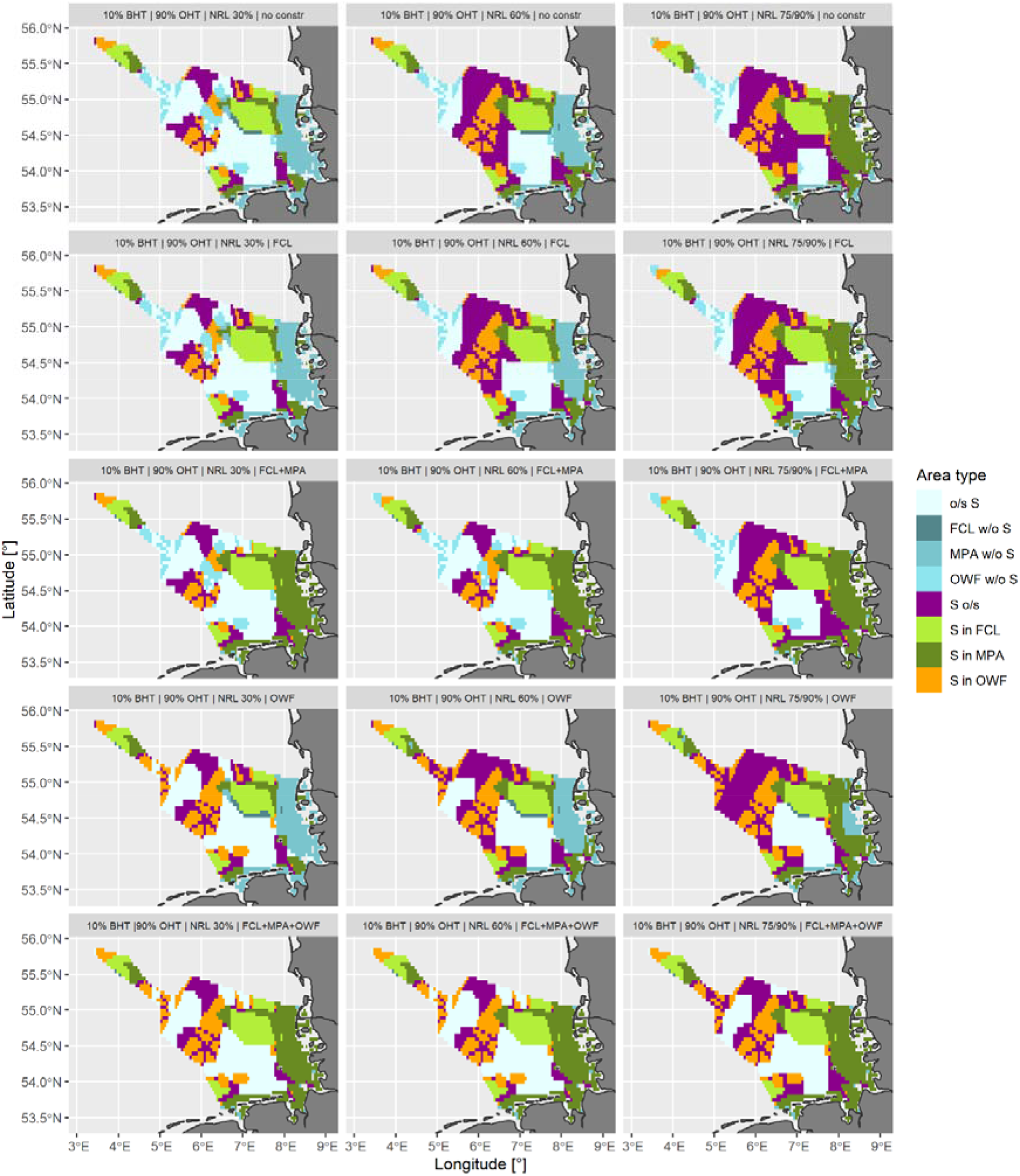
‘Prioritizr’-solutions for different for the different spatial scenarios and conservation targets.

**Figure 6.**
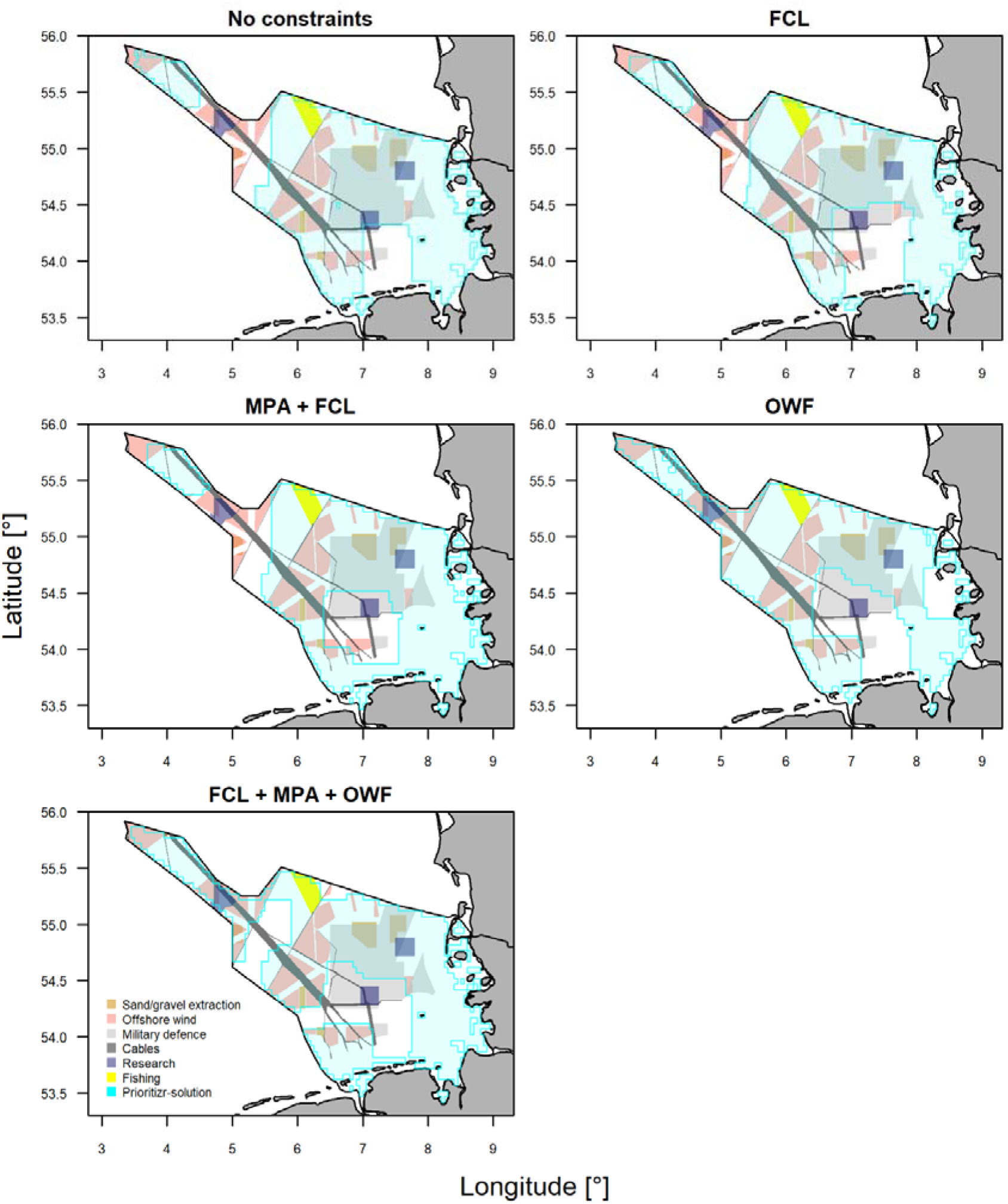
Overlap between “prioritizr”-solutions and areas for human activities under the maximum target scenarios (scenarios # 11-15, BHT= 10 %, OHT = 90 %, NRL1-5 = 90 %, NRLGroup7 = 75 %).

The analysis approach of this study uses the benthic habitat types as defined by the MSFD for the designation of NRL habitat groups. As a consequence, the overlap between corresponding MSFD habitat types and NRL habitat groups is high. While this might cause some redundancy in the inclusion of certain areas within the “prioritizr”-solutions, it allows for the evaluation of the targets for each directive/regulation. And even through the NRL habitat groups are based on the MSFD habitat types, they are not absolutely identical and hence warrant a separated consideration in the “prioritizr”-analysis.

So how could the spatial conflicts between the targets of the MSFD/NRL and the use of space for human activities be resolved? A part of the solution might be found in reducing the footprint of human activities on marine benthic habitats, e.g. by considering the conservation of original habitat types when planning OWF. For example, associated cable routes could be planned to favour the least length of cable, or foundations could be designed for permanent installation allow to re-power turbines without their removal. Further restrictions of trawl fishing while switching to passive and less invasive gears (Stelzenmüller et al., 2021) might further limit the impact of fisheries. Terrestrial sources for sand and gravel extraction or deposition might help to reduce impacts of coastal protection and harbour maintenance on marine benthic habitats. And artificial substrates such as concrete foundations for dikes and wind turbines might support the establishment of NRL group 2 & 3 habitats (macro algae forests & mussel beds), and it needs to be further discussed, if and how such artificial substrates might be included into the targets of the MSFD & NRL.

The pressure to use marine space will most likely increase, as demands for renewable energy, shipping, coastal protection or military defence will further increase under the scenario of global population growth and climate change (Li et al., 2023; Stelzenmüller et al., 2024). This is especially true for countries with limited access to marine space, which will try to accommodate as many activities as possible in waters under their jurisdiction. It remains therefore a most urgent matter of research on how the spatial demand of marine activities can be aligned with efficient solutions for marine conservation.

## Acknowledgements

This study is part of the Thünen-research initiative “Implementation of the Marine Strategy Framework Directive” (www.thuenen.de). Data from 4COffshore was purchased under the project “SeaUseTip” funded by the German Federal Ministry of Research, Education, Technology and Space (www.seausetip.de). The author thanks the German Agency for Nature Conservation (BfN) for the provision of benthic habitat data.

## Notes

### Competing Interest Statement

The authors have declared no competing interest.

