## Supplements for "A sea full of measures: EU conservation goals for benthic habitats will require wide-ranging spatial measures"

**Supplementary material**

**S1** R-code for analysing overlap between fisheries closures, MPA & OWF vs. MSFD benthic habitat types and NRL habitat groups using the package “prioritizr”.

\# nos.ger.area.rel: Raster stack with relative size of grid cells (as cost matrix)

\# ht.stk: Raster stack with habitat data

```
Library(prioritizr)

#### P1: How much habitat of each type is covered within fisheries closures? ----
p.ger.1<-
  problem(nos.ger.area.rel,ht.stk) %>%
  add_min_set_objective() %>%
  add_manual_targets(data.frame(feature = names(ht.stk),
                                type = "relative", sense = ">=",
                                target = 0)) %>%
  add_locked_in_constraints(fcl.ger) %>%
  add_binary_decisions() %>%
  add_cbc_solver()

s.ger.1<-solve(p.ger.1)

#### P2: How much habitat of each type is covered within MPA? ----

p.ger.2<-
  problem(nos.ger.area.rel,ht.stk) %>%
  add_min_set_objective() %>%
  add_manual_targets(data.frame(feature = names(ht.stk),
                                type = "relative", sense = ">=",
                                target = 0)) %>%
  add_locked_in_constraints(merge(fcl.ger,mpa.ger)) %>%
  add_binary_decisions() %>%
  add_cbc_solver()

s.ger.2<-solve(p.ger.2)
plot(s.ger.2)

#### P3: How much habitat of each type is covered within OWF? ----
p.ger.3<-
  problem(nos.ger.area.rel,ht.stk) %>%
  add_min_set_objective() %>%
  add_manual_targets(data.frame(feature = names(ht.stk),
                                type = "relative", sense = ">=",
                                target = 0)) %>%
  add_locked_in_constraints(owf.ger) %>%
  add_binary_decisions() %>%
  add_cbc_solver()

s.ger.3<-solve(p.ger.3)
plot(s.ger.3)

#### P4: How much habitat of each type is covered within MPA & OWF? ----
p.ger.4<-
  problem(nos.ger.area.rel,ht.stk) %>%
  add_min_set_objective() %>%
  add_manual_targets(data.frame(feature = names(ht.stk),
                                type = "relative", sense = ">=",
                                target = 0)) %>%
  add_locked_in_constraints(merge(fcl.ger,mpa.ger,owf.ger)) %>%
  add_binary_decisions() %>%
  add_cbc_solver()

s.ger.4<-solve(p.ger.4)
plot(s.ger.4)
```

**S2** R-code for implementing the spatial optimization with “prioritzr” for the MSFD & NRL targets as specified in Table 2 of the main manuscript.

\# ht.stk.a: Re-scaled (by factor rfc) raster stack with habitat data

\# spms: Re-scaled (by factor rfc) stack with constraint data (i.e. spatial measures)

```
# Define coverage targets
bht.tgt<-rep(0.1,3)
oht.tgt<-rep(0.9,3)
nrl.tgs<-c(0.3,0.6,0.9)
nrl.sdt<-c(0.3,0.6,0.75)


## Set prioritizr-prameters ----
cov.vec<-c(0.10,0.30,0.60,0.90)
type.vec<-c("no","FCL","MPA","MPA_OWF")
pnlty<-0.01
ef<-0.5  # Edge factor
rfc<-10  # Spatial rescaling factor to easy computational load
tl<-1800 # Define solver time out in sec

# Rescale cost matrix (relative area size)
area.new<-nos.ger.area.rel %>% aggregate(fact=rfc)
new.na.idx<-values(area.new) %>% is.na %>% which
new.v.idx<-values(area.new) %>% is.na %>% not %>% which
ref.rscld.area<-cellSize(area.new) %>% divide_by(1e6)
area.rscld<-ref.rscld.area/(ref.rscld.area[new.v.idx] %>%
  sum (na.rm=T))
values(area.rscld)[new.na.idx]<-NA

# Rescale habitats-stack
ht.stk.a<-ht.stk %>% aggregate(fact=rfc)

# Set progress bar and counte
ll<-length(nrl.tgs)*nlyr(spms)
pb<-txtProgressBar(min=0,max=ll,style=3)
cnt<-1

for (i in 1:length(nrl.tgs)) {
        for (j in 1:nlyr(spms)){

          # Elaborate target values
          r.trgts<-c(rep(bht.tgt[i],15),rep(oht.tgt[i],4),rep(nrl.tgs[i],3),nrl.sdt[i])

          # Define prioritizr-problem
          if(j==1){
            p.x<-problem(area.rscld,ht.stk.a) %>%
                add_min_set_objective() %>%
                add_relative_targets(r.trgts) %>%
                add_boundary_penalties(penalty=pnlty,edge_factor=ef) %>%
                add_binary_decisions()%>%
                add_cbc_solver(time_limit=tl)
          } else {
            p.x<-problem(area.rscld,ht.stk.a) %>%
              add_min_set_objective() %>%
              add_relative_targets(r.trgts) %>%
              add_boundary_penalties(penalty=pnlty,edge_factor=ef) %>%
              add_locked_in_constraints(spms[[j]]) %>%
              add_binary_decisions()  %>%
              add_cbc_solver(time_limit=tl)
          }

          # Solve prioritizr-problems
          s.x<-solve(p.x)

          # Store problems & solutions
          if(cnt==1) s.scn.stk<-s.x else s.scn.stk<-c(s.scn.stk,s.x)
          if(cnt==1) p.scn<-list(p.x) else p.scn<-c(p.scn,p.x)

          names(s.scn.stk)[cnt]<-paste0("s_",nrl.tgs[i],"_",names(spms)[j])
          names(p.scn)[cnt]<-paste0("p_",nrl.tgs[i],"_",names(spms)[j])

          cnt<-cnt+1
          setTxtProgressBar(pb,cnt)
}}
```

**S3** Parameter selection for “prioritizr”-scenarios 1-15

**Figure S3.1.** Ratio between area demand (i.e. cost) vs. average coverage (mean feature representation) using different values for the “prioritizr”-parameters edge factor, penalty and time limit for different coverage targets (10 % & 60 %). Any point above the dashed line represents a preferable deviation from a 1:1 coverage:cost ratio (i.e. more representation with less costs). Hence the parameters of edge factor = 0.5, penalty = 0.01 and time limit = 1.800 seconds was identified to provide the overall best performance for the 10%- and 60%-coverage scenarios.

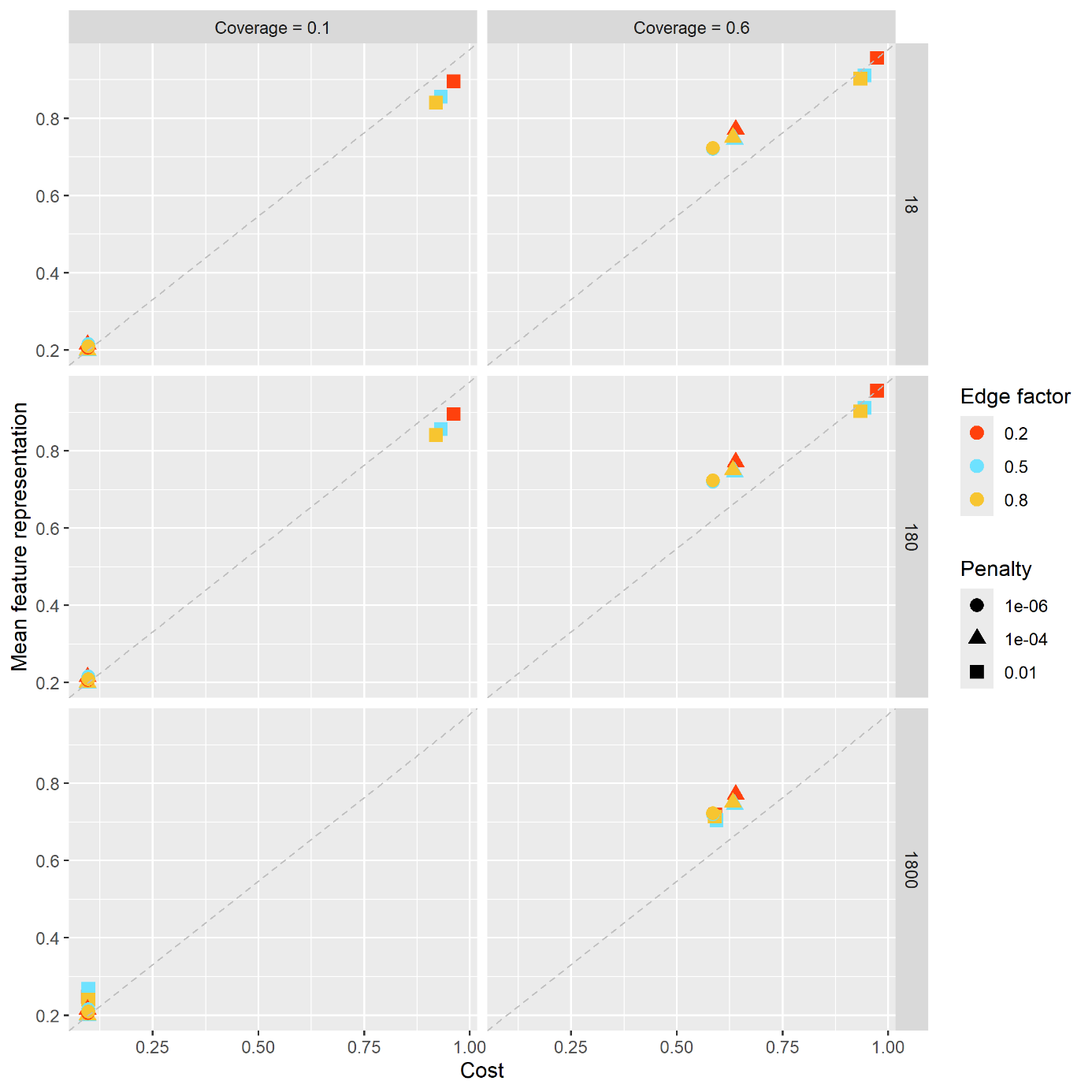

**Figure S3.2.** Maps of test runs with various parameter values. P=penalty [0.01, 1e-4, 1e-6], ef= edge factor [0.2, 0.5, 0.8], tl = time [18, 180, 1800 sec] limit and cvrg = coverage target [10 %, 60 %].

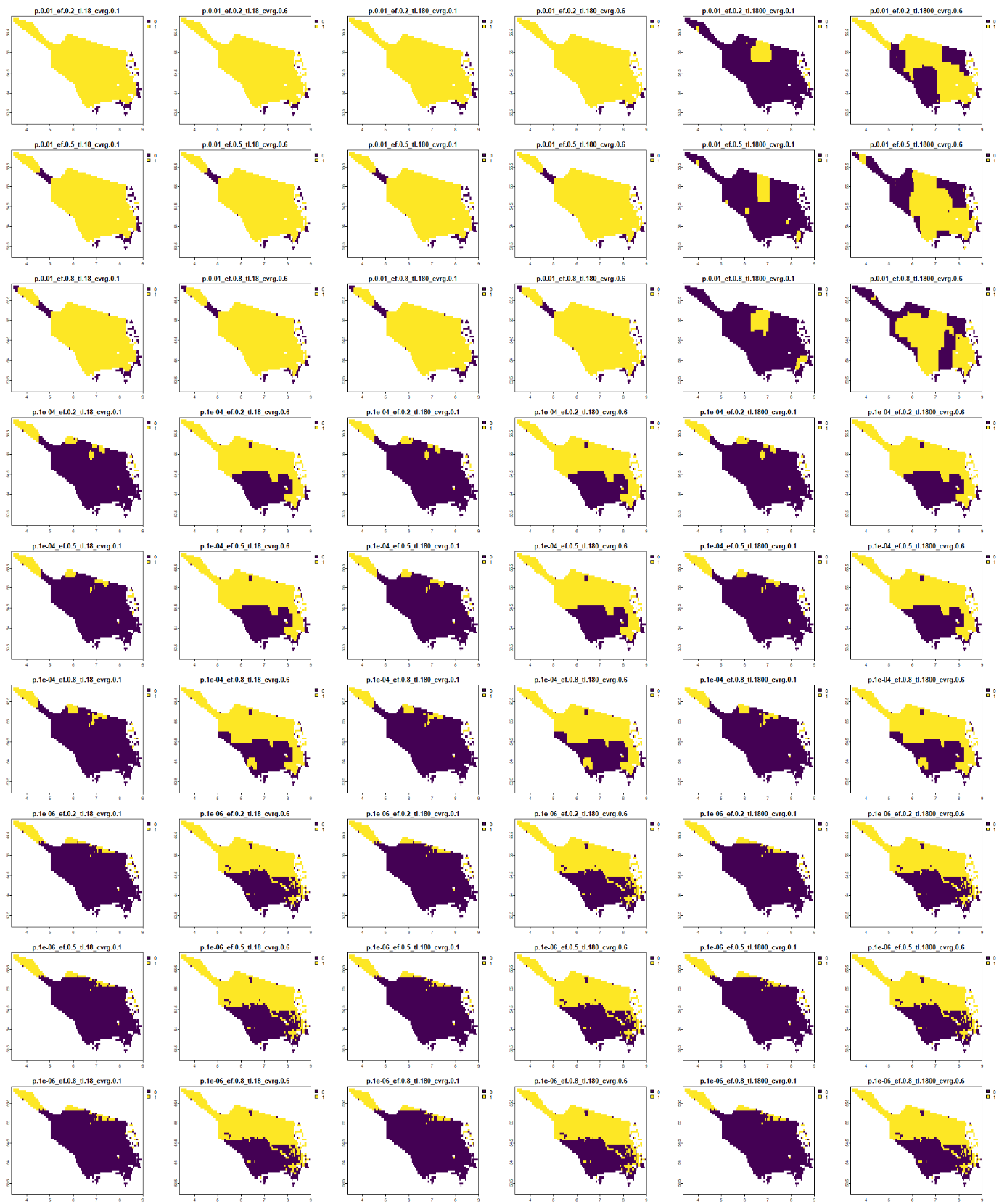
